# Neuroimaging alterations and relapse in early-stage psychosis

**DOI:** 10.1101/2021.12.10.472143

**Authors:** Marina Mihaljevic, Anisha Nagpal, Semra Etyemez, Zui Narita, Anna Ross, Rebecca Schaub, Nicola G. Cascella, Jennifer M. Coughlin, Gerald Nestadt, Frederik C. Nucifora, Thomas W. Sedlak, Koko Ishizuka, Vince Calhoun, Andreia V. Faria, Kun Yang, Akira Sawa

## Abstract

Recent reports have indicated that the occurrence of symptom exacerbation in early-stage psychosis could result in brain changes. Such a symptom exacerbation is frequently called relapse, which underlies a poorer disease outcome. Thus, it is important to identify neuroimaging alterations that are specifically seen in patients who experience relapse in early-stage psychosis. We hypothesized that this specific patient group may be more homogenous in disease-associated signatures likely to be linked to relapse, compared with the overall patient group. Such sub-stratified patient group and neuroimaging signatures would be useful for the biological understanding of relapse. To address this goal, we conducted a cross-sectional study (85 patients with early-stage psychosis and 94 healthy controls) with the use of medical records in a retrospective manner. To define the specific sub-group with the past experience of relapse, we used hospitalization due to psychotic symptom exacerbation, according to many publications that used this factor as a proxy for relapse. By examining resting-state functional connectivity (FC) for the study subjects, we validated our hypothesis and defined 131 FCs possibly associated with relapse. Through these studies, 3 brain regions (the thalamus, precentral gyrus, and dorsal anterior cingulate cortex) were underscored.

## INTRODUCTION

The trajectory of psychotic disorders after onset is heterogeneous, and many cases show deteriorating courses (Kahn et al., 2015, Owen et al., 2016). One of the major determinants of poorer prognosis is the occurrence of relapse, symptom exacerbations after a period of improvement (Suvisaari et al., 2018, Wunderink et al., 2020), which may be caused by intrinsic vulnerability but also by non-adherence to antipsychotic (AP) medication and co-morbid substance use disorder (Alvarez-Jimenez et al., 2012, Bowtell et al., 2018, Caseiro et al., 2012). Recent emerging evidence has consistently indicated that the occurrence of relapse in early-stage psychosis, regardless of reasons, could result in brain changes (Andreasen et al., 2013, Griffa et al., 2019, Rubio et al., 2022, Voineskos et al., 2020), which likely underlies the poorer disease outcome. For example, a pioneer study described that relapse duration was related to significant decreases in both total cerebral volume and frontal brain measures (Andreasen et al., 2013). This study implies that extended periods of relapse may have a negative effect on patients with schizophrenia, suggesting the importance of preventing and intervening with relapse. This notion is consistent with the observation of changes in structural connectivity among different stages of psychosis related to relapse in the staging model (Griffa et al., 2019), and the results of altered functional connectivity after relapse in patients with psychosis regardless proper AP maintenance (Rubio et al., 2022). Furthermore, a more recent paper reported adverse effects of relapse, independent of the effects of AP medications, on brain structure (Voineskos et al., 2020).

Taken together, it is important to identify neuroimaging alterations that are specifically seen in patients who experience relapse in early-stage psychosis. We hypothesized that this specific patient group, if defined in a proper manner, may be more homogenous in disease-associated signatures likely to be linked to relapse, compared with the overall patient group. Such sub-stratified patient group and signatures would be useful for a biological understanding of relapse. To address this goal, we used our cohort of early-stage psychosis (Coughlin et al., 2018, Faria et al., 2021, Kamath et al., 2018, Narita et al., 2021, Wang et al., 2019) and conducted a cross-sectional study with the use of medical records in a retrospective manner. We used hospitalization due to psychotic symptom exacerbation, as a simple and objective indicator of relapse, according to many publications that used this factor as a proxy for relapse (Addington et al., 2013, Csernansky et al., 2002, Olivares et al., 2013, Voineskos et al., 2020). By examining resting-state functional connectivity (FC) for the study subjects, we validated our hypothesis, defining FCs and specific brain reigons possibly associated with relapse.

## METHODS AND MATERIALS

### Study cohort

We recruited patients within 2 years after onset, and confirmed diagnoses of psychotic disorders using the Structural Clinical Interview for DSM-IV Patient Edition (SCID) (First et al., 1996). Exclusion criteria included a history of head trauma, nasal trauma, nasal surgery, neurologic disorder, cancer, viral infection, and reported history of intellectual disability. Additionally, participants with an estimated IQ below 70 on the Hopkins Adult Reading Test were excluded (Schretlen et al., 2009). Patients who reported active substance abuse or produced a urine drug screen positive for illicit substance use, except cannabis, were excluded. Individuals who were pregnant or taking anti-inflammatory agents or experiencing any other psychiatric conditions were also excluded. Healthy controls (HCs) who had a psychiatric diagnosis or a family history of schizophrenia-spectrum disorder were excluded. Positive and negative symptoms (presence, severity) of patients were evaluated with the Scale for the Assessment of Negative Symptoms (SANS) and the Assessment of Positive Symptoms (SAPS) (Andreasen, 1990). See our past publications using the same cohort for more detail (Coughlin et al., 2018, Faria et al., 2021, Kamath et al., 2018, Narita et al., 2021, Wang et al., 2019). This study was approved by the Johns Hopkins Medicine Institutional Review Boards and in accordance with the Code of Ethics of the World Medical Association (1964 Declaration of Helsinki). All study participants provided written informed consent. Parental consent and assent were obtained for all participants under the age of 18 years.

### Clinical characterization

Multiple board-certified psychiatrists assessed self-reports and the Johns Hopkins electronic medical database for the patients. Hospitalization due to psychotic symptom exacerbation was used as a simple and objective outcome indicator of relapse, according to many publications that supported the utility in a successful manner (Addington et al., 2013, Csernansky et al., 2002, Olivares et al., 2013, Voineskos et al., 2020). Accordingly, we used psychiatric hospitalization as a proxy for relapse. The Johns Hopkins electronic medical database and self-reports were used to identify patients who experienced psychiatric hospitalization(s) due to psychotic exacerbation (the relapse group: R group).

Medical records for treatment adherence, duration between hospitalizations, and cannabis use were examined, and four patients were excluded due to unclear medical history and hospitalization records. In addition, one patient was excluded due to head motion (see details in the “3-Tesla MRI brain imaging” section). Together, in this study, we investigated data from 94 HCs and 85 patients (31 in the R group). Patients were diagnosed as schizophrenia (n = 43), schizoaffective disorder (n = 14), schizophreniform disorder (n = 2), bipolar disorder with psychotic features (n = 19), major depressive disorder with psychotic features (n = 5), and not otherwise specified psychotic disorder (n = 2).

Chlorpromazine (CPZ) equivalent doses for antipsychotics using the defined daily dose method (Leucht et al., 2016). Duration of illness (DOI) [between the onset and resting state functional Magnetic Resonance Imaging (rs-fMRI) scan] and antipsychotic doses were obtained through self-reports and confirmed through the Johns Hopkins electronic medical database. In the patient group, 5 patients were treated with first-generation antipsychotics, 67 with second-generation antipsychotics, and 6 with both first and second-generation antipsychotics. Seven patients were unmedicated.

### 3-Tesla MRI brain imaging

Study participants were scanned using a Phillips 3T MRI scanner. Details about data acquisition and processing have been reported in our previous publications (Faria et al., 2021, Mori et al., 2016). Specifically, the rs-fMRI data were obtained with the following parameters: axial orientation, original matrix 80×80, 36 slices, voxel size 3×3×4 mm, TR/TE 2000/30 ms, 210 time points.

MRICloud (www.MRICloud.org), a cloud-based high-throughput neuroinformatic platform developed by one of the co-authors (A.F.) of the present study and other researchers at Johns Hopkins University, was used for automatic whole brain segmentation. In this study, the segmentation was based on a multi-atlas library derived from multiple healthy young adults (Mori et al., 2016, Wu et al., 2016). The atlas and its 3D visualization are available at MRICloud BrainKnowledge: https://brainknowledge.anatomyworks.org. The segmentation process includes skull stripping, sequential linear and diffeomorphic mapping, and multi-atlas labeling by joint label fusion (Tang et al., 2013). The rs-fMRI data were then processed using MRICloud as follows: images were slice-time-corrected to adjust for differences in the acquisition time between slices, realigned to the first image using rigid body registration to adjust for motion, and then co-registered to the MPRAGE. COMPCOR, an approach that uses signals from the deep white matter and ventricles to estimate these artifacts and then remove them from grey matter time courses, was used to correct signals from non-neuronal physiological activity, such as respiration and cardiac pulsation. Seed-by-seed correlation matrices were obtained from the nuisance-corrected time courses and z-transformed by Fisher’s method. At last, we obtained 3,003 pairwise resting-state z-correlations between 78 brain regions of interest (ROIs).

While realignment and head motion evaluation were preprocessed as mentioned above, we further calculated framewise displacement (FD) using the six motion parameters matrix of each subject (Power et al., 2012) to estimate head motion. Study participants with FD larger than 0.3 were considered outliers and excluded from analysis (Achterberg and van der Meulen, 2019, DeSerisy et al., 2020, Li et al., 2017). Accordingly, one subject was identified as an outlier and excluded from the analyses.

### Statistical analysis

R 3.5.1 was used to perform statistical analysis. T-test and chi-squared test were conducted to compare demographics between groups for continuous and categorical variables, respectively.

ANCOVA (analysis of covariance) with age, sex, race, handedness, smoking status, and FD as covariates was conducted to compare FCs between the entire patient group and HC, and between the R group and HC. Correlation analysis was further conducted to evaluate the effects of CPZ dose and DOI in significant results. The Benjamini-Hochberg procedure was performed for multiple comparison correction for all the analyses where multiple statistical tests were involved (Glickman et al., 2014). Adjusted p-values were presented as q-values. Results with q-values smaller than 0.05 were considered significant.

## RESULTS

The study cohort consisted of 94 HCs and 85 patients. Thirty-one patients had experienced relapses, and we defined this subset as the R group. The entire patient group had more males and more cigarette smokers compared to HCs (**Table 1**). Similarly, the R group had more males and more cigarette smokers compared to HCs (**Table 1**). All demographic factors were adjusted in the data analysis.

**Table 1.** Main characteristics of participants. Abbreviations: HC, healthy control; R, patients who experienced relapse

|  | HC<br>(n=94) | Entire patient<br>group<br>(n=85) | R group<br>(n=31) | HC vs<br>Entire patient<br>group<br>p-value | HC vs<br>R group<br>p-value |
| --- | --- | --- | --- | --- | --- |
| Age<br>(mean±SD) | 23.5±3.8 | 22.3±4.6 | 22.4±4.2 | 0.078 | 0.211 |
| Sex<br>(male %) | 44.7 | 71.8 | 71.0 | **<0.001** | **0.011** |
| Race<br>(white %) | 37.2 | 49.4 | 35.5 | 0.100 | 0.861 |
| Smoking<br>(yes %) | 4.3 | 27.1 | 32.3 | **<0.001** | **<0.001** |
| Handedness<br>(right %) | 89.4 | 89.4 | 80.6 | 0.991 | 0.208 |
| Note: p-value <0.05 is presented in bold |  |  |  |  |  |

Compared to HC, we identified 60 and 131 significant FCs in the entire patient group and the R group, respectively, after multiple comparison correction (**Figure 1, Table S1**). We further conducted correlation analysis to evaluate the effects of antipsychotic medication and DOI on the significant results and didn’t find any significant correlations, which suggested that the differences between the R group and HC may not be caused by antipsychotics and DOI. Among these 60 FCs with significant alterations in the patient group, 51 of them were also significantly altered in the R group, implying that the R group may be the main contributor to the changes observed in the entire patient group.

**Figure 1.**
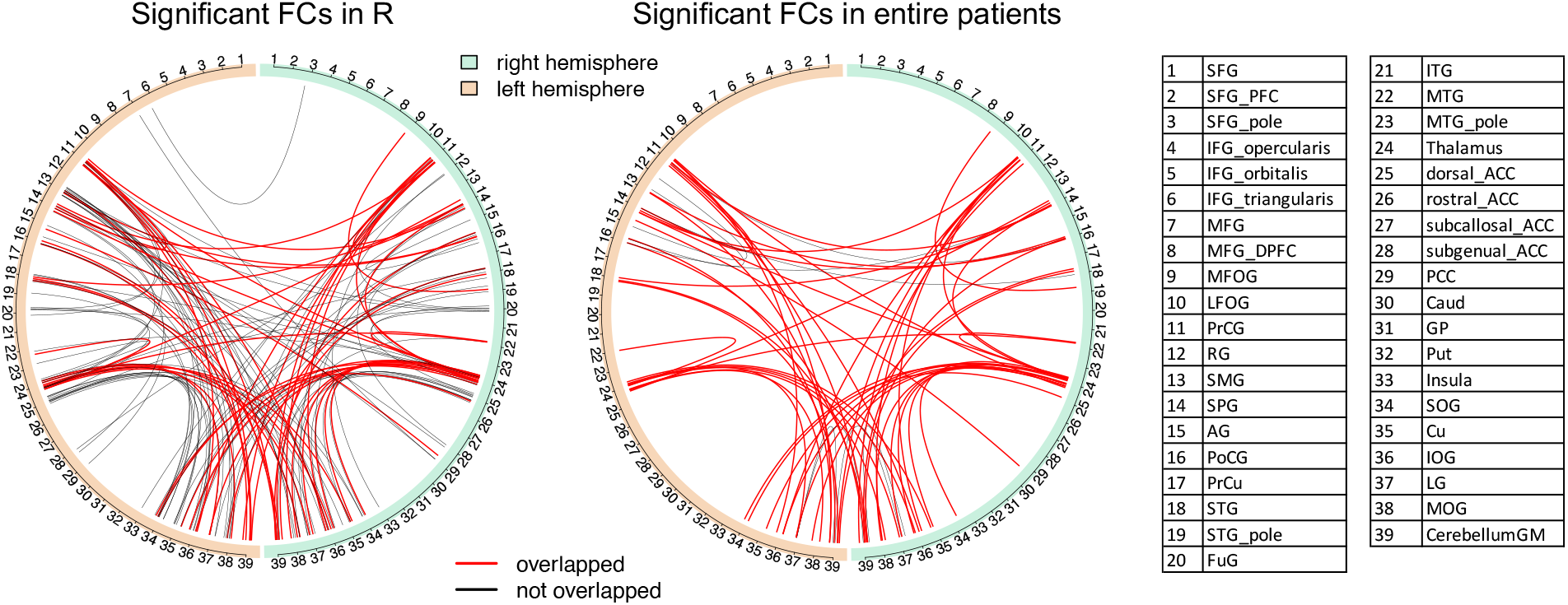
Chord diagram of significant brain functional connectivity (FC). We identified 131 and 60 significant FCs after multiple comparison corrections (q-value <0.05) between patients who experienced relapse (the R group) and healthy controls (HC) and between the entire patient group and HC, respectively. Red lines represent those 51 significant FCs with significant changes in both the R group and the entire patient group. Black lines represent other significant FCs with changes only in the R group or only in the entire patient group. The full name for the abbreviations of regions of interests (ROIs) used in this figure is listed in **Table S1**.

We further compared the effect sizes and variations of the 51 FCs commonly altered in both the entire patient and R groups to test whether the R group is more homogeneous than the entire patient group. Interestingly, we found that the effect sizes of alterations in those 51 FCs in the R group were significantly larger than the effect sizes in the entire patient group (mean in the R group: 0.895; mean in the entire patient: 0.637; p-value = 5.649e-26). Meanwhile, the R group had significantly smaller variations in those 51 FCs compared to the entire patient group (mean in the R group: 0.075; mean in the entire patient group: 0.082; p-value = 4.995e-02). These results showed that the R group is more homogeneous than the entire patient group. Studying the R group may be useful for us to study biological mechanisms and biomarkers associated with relapse, in particular those as the devastating outcome of relapase.

Accordingly, we further examined brain regions that participated in 131 significant FCs in the R group compared to HCs. Thalamus, precentral gyrus, and dorsal anterior cingulate cortex are the top 3 regions with the most number of significant FCs in the R group (**Figure 2**).

**Figure 2.**
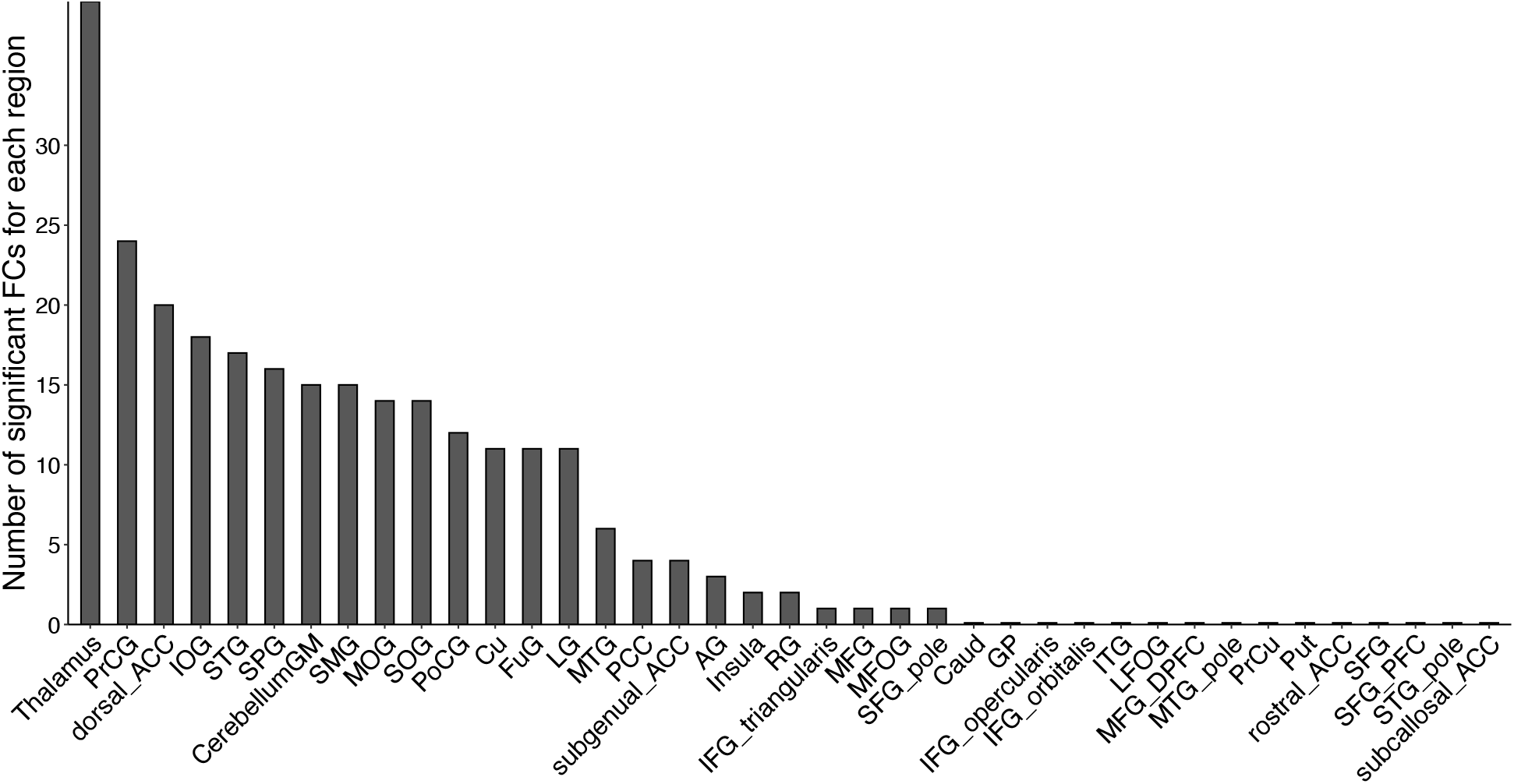
Bar plot of the number of significant brain functional connectivity (FC) linked to each brain regions of interest (ROI) We identified 131 significant FCs between patients who experienced relapse (R) and healthy controls (HC). None of these FCs was correlated with antipsychotics and duration of illness. The top 3 ROIs accounting for 64% of significant FCs in the R group were thalamus, precentral gyrus (prCG), and dorsal anterior cingulate cortex (ACC). The full name for the abbreviations of ROIs used in this figure is listed in **Table S1**.

## DISCUSSION

The goal of the present study is to look for an effective strategy to study biological mechanisms associated with relapse, particularly those underlying the devastating outcome of relapse, which is a key determinant of disease trajectory for patients with early-stage psychosis. Thus, it is important to identify neuroimaging alterations that are more specifically seen in patients who experience relapse in early-stage psychosis. We hypothesized that this specific patient group may be more homogenous in disease-associated signatures likely to be linked to relapse, compared with the overall patient group. To define the specific sub-group with the past experience of relapse, we used hospitalization due to psychotic symptom exacerbation, according to many publications that used this factor as a proxy for relapse. By examining resting-state FCs for the study subjects, we validated our hypothesis and defined 131 FCs and 3 brain regions possibly associated with relapse.

Relapse is a complex condition to which many factors, such as AP medication cessation, substance abuse, and stressful life events, may contribute (Taylor and Jauhar, 2019). Thus, longitudinal studies are expected to address the cause and impact of relapse in a precise manner. Meanwhile, a systematic review regarding relapse in schizophrenia reported that hospitalization was the most widely used factor as a proxy for relapse by representing symptom exacerbation (62% of publications) (Olivares et al., 2013).

The merit of using hospitalization is that it is objective, scalable, quantitative, and good as a proxy for large multi-institutional studies, including those in a cross-sectional design (Addington et al., 2013). We acknowledge potential caveats in interpreting hospitalization as an endpoint for symptom exacerbation, because other factors such as suicidality and family pressure may play a role. At least in the present study, we aimed to minimize this concern by using an electronic medical database effectively. Taken all together, in this cross-sectional study that is expected to be a prototype for scalable and multi-institutional projects (including data from multiple countries) in near future, we used the proxy of relapse to study brain alternations specifically found in a subgroup of patients in early-stage psychosis who experienced relapse. In parallel, a prospective study design may be able to more ideally answer questions about whether brain alteration in patients who experienced relapse could reflect intrinsic vulnerability to relapse or brain changes elicited by relapse.

## ACKNOWLEDGMENTS AND DISCLOSURES

This study is supported by NIH grants MH-092443, MH-094268, MH-105660, and MH-107730; foundation grants from Stanley and RUSK/S-R (to AS), an award from Brain and Behavior Research Foundation (to KY), and a Fulbright fellowship (to MM). Study recruitment was in part funded by Mitsubishi Tanabe Pharma Corporation. The authors thank Ms. Yukiko Y Lema for organizing the manuscript.

All authors have nothing to disclose.

