## Supplementary table for "Neuroimaging alterations and relapse in early-stage psychosis"

Postal address: 600 N Wolfe ST, Baltimore, MD 21287, USA

**Keywords:** relapse, early-stage psychosis, resting-state functional MRI, large-scale brain network, dorsal anterior cingulate cortex, thalamus.

**Table S1. Abbreviations and full names of brain regions of interest**

| Abbreviation | Full Name |
|---|---|
| SFG | Superior frontal gyrus |
| SFG pole | Superior frontal gyral pole |
| SFG_PFC | Superior frontal gyrus_prefrontal cortex |
| MFG | Middle frontal gyrus |
| MFG_DPFC | Dorsal prefrontal pars of middle frontal cortex |
| MFOG | Middle fronto-orbital gyrus |
| IFOG | Lateral fronto-orbital gyrus |
| IFG_opercularis | Inferior frontal gyrus (pars opercularis) |
| IFG_orbitalis | Inferior frontal gyrus (pars orbitralis) |
| IFG_triangularis | Inferior frontal gyrus (pars triangularis) |
| RG | Gyrus rectus |
| PrCG | Precentral gyrus |
| PoCG | Postcentral gyrus |
| SPG | Superior parietal gyrus |
| SMG | Supramarginal gyrus |
| AG | Angular gyrus |
| PrCu | Pre-cuneus |
| Cu | Cuneus |
| LG | Lingual gyrus |
| SOG | Superior occipital gyrus |
| IOG | Inferior occipital gyrus |
| FuG | Fusiform gyrus |
| MOG | Middle occipital gyrus |
| STG | Superior temporal gyrus |
| STG pole | Superior temporal gyral pole |
| MTG | Middle temporal gyrus |
| MTG pole | Middle temporal gyrus pole |
| ITG | Inferior temporal gyrus |
| PCC | Posterior cingulate gyrus |
| Insula | Insula |
| Caud | Caudate nucleus |
| GP | Globus pallidus |
| Put | Putamen |
| Thalamus | Thalamus |
| dorsal_ACC | Dostral anterior cingulate gyrus |
| rostral_ACC | Rorsal anterior cingulate gyrus |
| subcallosal_ACC | Subcallosal anterior cingulate gyrus |
| subgenual_ACC | Subgenual anterior cingulate gyrus |
| CerebellumGM | Cerebellum gray matter |
